# Transcriptomic investigation of the molecular mechanisms underlying resistance to the neonicotinoid thiamethoxam and the pyrethroid lambda-cyhalothrin in *Euschistus heros* (Hemiptera: Pentatomidae)

**DOI:** 10.1101/2023.05.09.539981

**Authors:** Ewerton C. Lira, Antonio R.B. Nascimento, Chris Bass, Celso Omoto, Fernando L. Cônsoli

## Abstract

Strains of *Euschistus heros* (Hemiptera: Pentatomidae) with resistance to thiamethoxam (NEO) and lambda-cyhalothrin (PYR), generated by selection with these insecticides in the laboratory, have been recently reported in Brazil. However, the mechanisms conferring resistance to these insecticides in *E. heros* remain unresolved. We utilized comparative transcriptome profiling and single nucleotide polymorphism (SNP) variant calling of susceptible and laboratory-selected resistant strains of *E. heros* to investigate the molecular mechanism(s) underlying resistance. The *E. heros* transcriptome was assembled using approximately 190.1 million paired-end reads, which generated 91,673 transcripts with a mean length of 720 bp and N50 of 1795 bp. Approximately, 54.8% of the assembled transcripts ware functionally annotated against the NCBI *nr* database, with most sequences (43%) being similar to the pentatomids *Halyomorpha halys* (43%) and *Nezara viridula* (29%). Comparative gene expression analysis between the susceptible (SUS) and NEO strains identified 215 significantly differentially expressed (DE) transcripts. DE transcripts associated with the metabolism of xenobiotics were all up-regulated in the NEO strain. The comparative analysis of the SUS and PYR strains identified 204 DE transcripts, including an esterase (esterase FE4), a glutathione-S-transferase, an ABC transporter (ABCC1), and aquaporins that were up-regulated in the PYR strain. We identified 9,588 and 15,043 non-synonymous SNPs in the PYR and NEO strains respectively in comparisons with the SUS strain. One of the variants (D70N) detected in the NEO strain occurs in a subunit (α5) of the nicotinic acetylcholine receptor, the target-site of neonicotinoid insecticides. Nevertheless, the position of this residue was found very variable among α5 from insect species. In conclusion, neonicotinoid and pyrethroid resistance in laboratory-selected strains of *E. heros* is associated with a potential metabolic resistance mechanism mediated by the overexpression of several proteins commonly involved in the three phases of xenobiotic metabolism. Together these findings provide insight into the potential basis of resistance in *E. heros* and will inform the development and implementation of resistance management strategies against this important pest.

**Highlights:**

- 419 DE genes were observed in *E. heros* insecticide-resistant strains
- 24,631 SNPs were identified in *E. heros* insecticide-resistant strains
- *E. heros* insecticide-resistant strains overexpress metabolic resistance genes
- Lambda-cyhalothrin-resistant *E. heros* overexpresses cuticular proteins
- Thiamethoxam-resistant *E. heros* carries the target-site mutation D70N in nAChRalpha5

## 1. Introduction

Neonicotinoids are nicotinic acetylcholine receptor (nAChR) agonists that maintain the receptor in a continuously activated state by affecting the dynamic ratio of Na^+^ and K^+^, which consequently leads to insect death by hyperexcitation of the nervous system (Ihara et al., 2017; Seifert, 2014; Tomizawa and Casida, 2005). Pyrethroids are sodium channel modulators that alter the normal function of insect nerves by modifying the kinetics of voltage-gated sodium channels (VGSCs). Pyrethroids can have VGSC state binding preferences, but many pyrethroids preferentially bind to open sodium channels, stabilize the open state, and subject neurons to prolonged activation of the membrane electrical potential (Silver et al., 2014; Soderlund, 2010).

Neonicotinoids and pyrethroids have been used on a large scale over extensive areas to control the neotropical brown stink bug *Euschistus heros* (F.) (Hemiptera: Pentatomidae), one of the most important insect pests of soybean crops in Brazil (Sosa[Gómez et al., 2020). Over the last decade *E. heros* has gained prominence, becoming the major pest targeted for chemical control, particularly after the release of Bt-soybean for management of lepidopteran pests of soybean (Panizzi et al., 2022). Currently, according to the Phytosanitary Pesticide System there are 86 pesticides registered in Brazil, including simple formulations and mixtures of active ingredients for the control of *E. heros* in soybean crops (Agrofit, 2023). However, these pesticides belong predominantly to only three insecticide classes organophosphates, pyrethroids, and neonicotinoids encompassing just three distinct modes of action. As a result, several authors have reported a decrease in susceptibility to different pesticides in populations of *E. heros* spread over extensive areas in Brazil. Such decrease in the frequency of the susceptible phenotypes in field conditions has facilitated the selection of two strains of *E. heros* with resistance to the neonicotinoid thiamethoxam or to the pyrethroid lambda-cyhalothrin (Castellanos et al., 2019; Somavilla et al., 2020a; Somavilla et al., 2020b; Sosa[Gómez et al., 2020; Sosa[Gómez et al., 2001; Sosa-Gómez and Silva, 2010; Tibola et al., 2021; Tuelher et al., 2018). The ease with which resistance can be selected in the laboratory serves as a warning of the high risk of field evolved resistance in *E. heros* to neonicotinoids and pyrethroids if integrated pest management and insect resistance management strategies are not employed.

Target site mutations and enhanced metabolism are common resistance mechanisms of insects to neonicotinoids (Sahani et al., 2022) and pyrethroids (Keïta et al., 2021; Liu, 2012). Even though neonicotinoids and pyrethroids are old molecules and the physiological and/or molecular mechanisms of insect resistance have been reported for numerous insect species, little information on the molecular mechanisms of insecticide resistance in stinkbugs is available, in particular for *E. heros* (King et al., 2023; Singh et al., 2023). The use of omics approaches in the last decade has advanced our understanding of the molecular genetics of insecticide resistance in insects, and allowed the identification of complementary, multiple mechanisms of resistance existing in a single genome (ffrench-Constant, 2013).

In this study we used a transcriptomic approach to investigate the molecular basis of *E. heros* resistance to thiamethoxam and lambda-cyhalothrin. We characterized the transcriptome of susceptible and thiamethoxam and lambda-cyhalothrin resistant strains of *E. heros,* compared the gene expression patterns between susceptible and resistant strains, and identified single nucleotide polymorphisms in transcripts from each insecticide-resistant strain when compared to the susceptible strain of *E. heros.* Identification of putative molecular mechanism(s) involved in the resistance of lab-selected thiamethoxam and lambda-cyhalothrin resistant strains of the neotropical brown stink bug is a first step towards the development of tools for large-scale resistance monitoring to support the pest management decision process and the development and implementation of resistance management strategies.

## 2. Material and Methods

### 2.1. Insect strains

Adults from three strains of *E. heros* were used for sequencing: a susceptible reference strain (SUS) collected in soybean fields in the field station at Luiz de Queiroz College of Agriculture, University of São Paulo, Piracicaba, SP, Brazil and two laboratory-selected resistant strains, one to the pyrethroid lambda-cyhalothrin (PYR) and the other to the neonicotinoid thiamethoxam (NEO) obtained from a field population collected in Luís Eduardo Magalhães – BA and Londrina - PR (Tibola et al., 2021). The SUS strain has been maintained in the absence of selection pressure for more than 6 years under laboratory conditions and was donated by the company Pragas.com® (Piracicaba, São Paulo State, Brazil). The resistant strains PYR and NEO were obtained by mass selection to the selected insecticides under laboratory conditions. The resistance ratio of PYR is 66.3-fold, while that of NEO is 44.3-fold (Tibola et al., 2021).

### 2.2. RNA extraction, library preparation and sequencing

Total RNA was isolated from four biological replicates, with each replicate representing a pool of ten adults of *E. heros* for each strain (SUS, PYR and NEO). RNA extraction was performed using the RNeasy Kit (Qiagen), according to the manufacturer’s guidelines. Genomic RNA was evaluated quantitatively in a Qubit fluorometer (Thermo Fisher Scientific, USA), and RNA integrity was verified by standard agarose gel electrophoresis (Sambrook and Russell, 2001) and RNA integrity number (RIN) determination in an Agilent Bioanalyzer. RNA samples were sent to the Functional Genomics Applied to Agriculture and Agri-energy Instrumentation Center (USP/ESALQ, Piracicaba, SP) for cDNA library preparation and paired-end (PE) (2×100 bp) Illumina sequencing on a NextSeq 550 platform.

### 2.3. De novo assembly and functional annotation

The raw reads obtained were assessed using FastQC (Andrews, 2010). Adapter sequences were trimmed, and low-quality bases (Phred quality score <20 bp) and short reads (< 25 bp) removed using Trimmomatic v0.39 (Bolger et al., 2014). The high-quality clean reads obtained from all libraries (susceptible and resistant) were used for the *de novo* assembly of a reference transcriptome of *E. heros* using Trinity v2.14.0 (https://github.com/trinityrnaseq/trinityrnaseq/wiki). The *de novo* transcriptome assembly was obtained using a *k-mer* size of 25 bp and a minimum transcript length of 300 bp. The reference *de novo* transcriptome was filtered before comparative differential expression analysis, and only transcripts with *transcript per million* (TPM) values ≥ 1 were further analyzed.

The filtered assembled transcripts were interrogated against the NCBI non-redundant protein database (nr-NCBI) using the BLASTX algorithm in Diamond (Buchfink et al., 2015), with an *e*-value cut-off of 10^-3^. The candidate protein-coding regions within assembled transcripts were extracted from the reference assembly using TransDecoder (https://transdecoder.github.io/) (Haas et al., 2013). The protein sequences predicted from candidate protein-coding regions were subjected to BLASTp search (*e*-value cut-off of 10^-3^) and were functionally characterized based on sequence ortholog assignments available in the EggNOG database (https://github.com/eggnogdb/eggnog-mapper) using EggNOG-mapper (Huerta-Cepas et al., 2019). EggNOG-mapper offers functional annotation against several databases, such as Pfam, Kyoto Encyclopedia of Genes and Genomes (KEGG) Pathway Database, Gene Ontology, and Clusters of Orthologous Genes (COG) database (Cantalapiedra et al., 2021).

### 2.4. Identification of Single Nucleotide Polymorphism (SNP)

For SNP identification was following the workflow for identifying short variants in RNA-seq data (https://gatk.broadinstitute.org/hc/en-us/articles/360035531192-RNAseq-short-variant-discovery-SNPs-Indels). The high-quality reads obtained from all libraries (susceptible and resistant) were aligned against the *de novo* transcriptome using the *bwa-mem* algorithm in the BWA Aligner software using default parameters (Li and Durbin, 2009). SAMtools was used to convert and group alignment results to BAM files according to the treatment (SUS, NEO and PYR) (Li et al., 2009). Reads aligned and grouped were ordered by coordinates using the *AddOrReplaceReadGroups* tool (Picard Toolkit, https://broadinstitute.github.io/picard). Duplicate reads were removed using the *MarkDuplicates* tool (Picard Toolkit) and alignments that cover introns were reformatted using the *SplitNCigarReads* tool, available in the Genome Analysis Toolkit (GATK) v. 42.6.1 (https://gatk.broadinstitute.org/hc/en-us). The *HaplotypeCaller* tool included in the package GATK was used in SNP calling with the following arguments: --*standard-min-confidence-threshold-for-calling* of 30 and *--min-base-quality-score* of 25. The initial list of identified variants was filtered using the *SelectVariants* tool (GATK) with the *--select-type-to-include* argument to obtain a subset containing only SNPs. The lists of SNPs from each treatment (SUS, NEO and PYR) were filtered with a series of quality metrics [depth of coverage (DP) < 10, quality by depth (QD) < 5, quality of mapping (MQ) < 40, and the Phred-scaled probability that there is strand bias at the site (FS) > 60] using *VariantFiltration* tool (GATK). As our main objective was to characterize SNPs that could explain the resistance of *E. heros* to thiamethoxan and/or lambda-cyhalothrin, SNPs identified in the resistant strains were compared with the reference susceptible strain and their effects were verified using the *SnpEff* software (Cingolani et al., 2013). SNPs within predicted coding regions were classified mainly as non-synonymous (missense variant) or synonymous. The *SnpSift* software (Cingolani et al., 2012) was used to select for non-synonymous SNPs. Selected transcripts with non-synonymous SNPs were functionally annotated by combining the list of SNPs with the functional annotation of the *E. heros* reference transcriptome.

### 2.5. Gene expression analysis

The determination of the abundance of transcripts in each library (SUS, PYR, and NEO) was performed by mapping the filtered reads against the reference *de novo* transcriptome assembly using RSEM with default parameters (Li and Dewey, 2011). The read counts obtained for transcripts were normalized using the *transcript per million* (TPM) method for RNA-Seq analysis (Abrams et al., 2019). TPMs values were then used for comparative analysis of transcript abundance (= differential expression - DE) between the susceptible (SUS) and each of the resistant strains (PYR and NEO) of *E. heros* using the DEseq2 package (Love et al., 2014). The resulting *p-values* in comparisons of expression levels were adjusted to avoid type I errors (false positives) using the Benjamini-Hochberg FDR method (Haynes, 2013), and only transcripts with a *log_2_* fold change (FC) ≥ |2| and a false discovery rate (FDR) p-adj ≤ 0.05 were considered differentially expressed. The differentially expressed transcripts were concatenated with the functional annotation data, and further subjected to functional analysis and GO terms functional enrichment analysis (Kolmogorov-Smirnov, *p* < 0.05) using the TopGO package (Alexa and Rahnenfuhrer, 2022).

## 3. Results

### 3.1. RNA-sequencing and de novo assembly

Sequencing of susceptible and resistant strains of *E. heros* yielded 213.9 million paired-end raw reads, ranging from 17 to 20.9 million reads per library (Table S1). Approximately 11% of raw reads were removed after read trimming and quality filtering, and 190.1 million (∼ 89%) of high-quality paired-end reads were used to assemble the *de novo* reference transcriptome of *E. heros*. The GC percent content of samples ranged from 39% to 41%, and the mapped ratio of reads against the *de novo* reference assembly was higher than 80% for all samples (Table S1). The *de novo* assembled transcriptome resulted in 131,025 transcripts and 79,808 genes, with a transcript mean length of 657 bp and an N50 of 1,638 bp. After filtering out transcripts with TPM values below 1, the final transcriptome contained 91,673 transcripts and 55,510 genes, with a transcript mean length of 720 bp and an N50 of 1,795 bp (Table S1).

### 3.2. Functional annotation

The search for assembled transcripts (BLASTx) and their predicted coding regions (CDS) (BLASTp) against the NCBI *nr* database successfully annotated 50,235 (54.8%) out of the 91,673 transcripts of the reference transcriptome (Fig. 1A), with most sequences (43%) matching those of the brown marmorated stink bug *Halyomorpha halys* (Stal) and *Nezara viridula* (L.) (29%) (Fig. 1B).

**Fig. 1.**
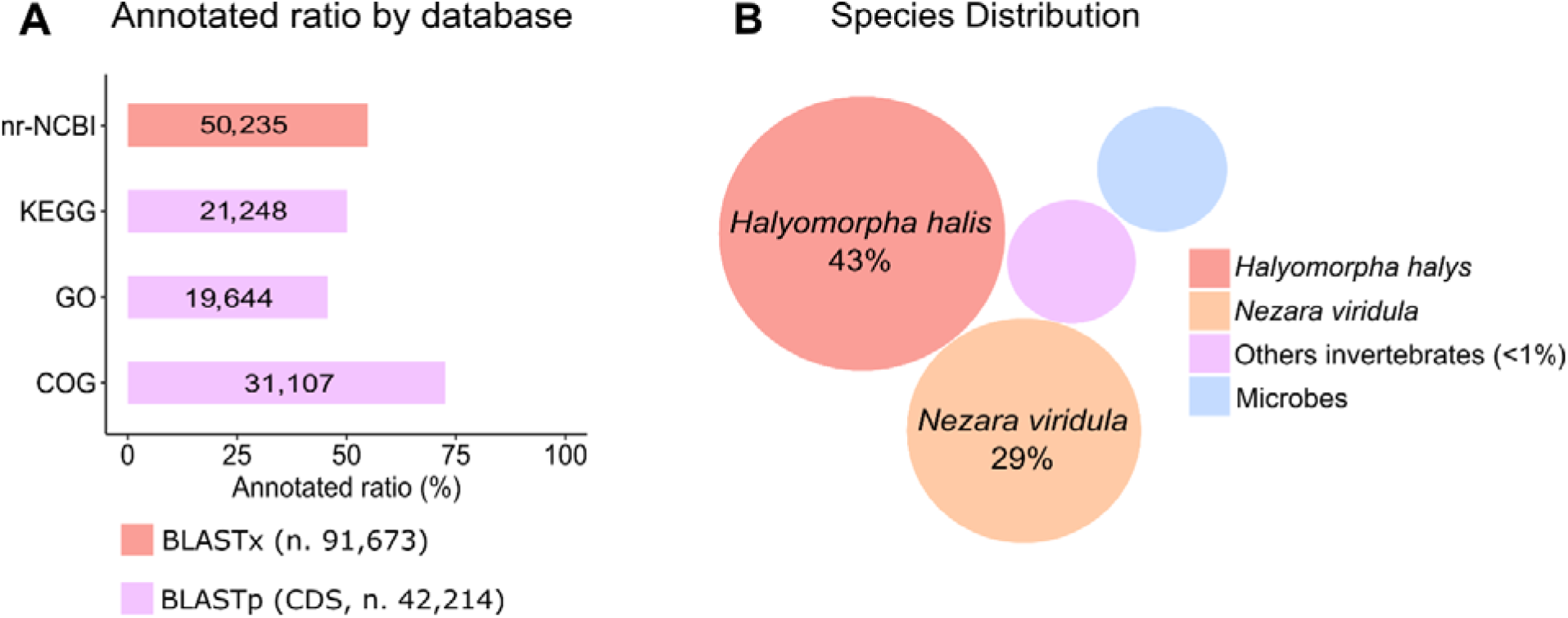
**(A)** Summary of the functional annotation of predicted proteins from the *de novo* transcriptome of *Euschistus heros* against different databases, and **(B)** species classification of transcripts from the *de novo* transcriptome of *E. heros* based on the BLAST results against the NCBI *nr* database.

Analysis of the functional classification of the 42,214 protein-coding regions identified using TransDecoder assigned gene ontology (GO) to 19,644 (46.53%) sequences, distributed in 73 functional groups divided in *biological processes* (32), *cellular components* (20), and *molecular functions* (21) categories (Fig. 2). The most representative GO terms in *biological processes* were related to *cellular process* (GO:0009987), *metabolic process* (GO:0008152) and *biological regulation* (GO:0065007); in *cellular components* were *cell part* (GO:0044464) and *cell* (GO: 0005623); and *binding* (GO: 0005488) and *catalytic activities* (GO: 0003824) in *molecular functions* (Fig. 2).

**Fig. 2.**
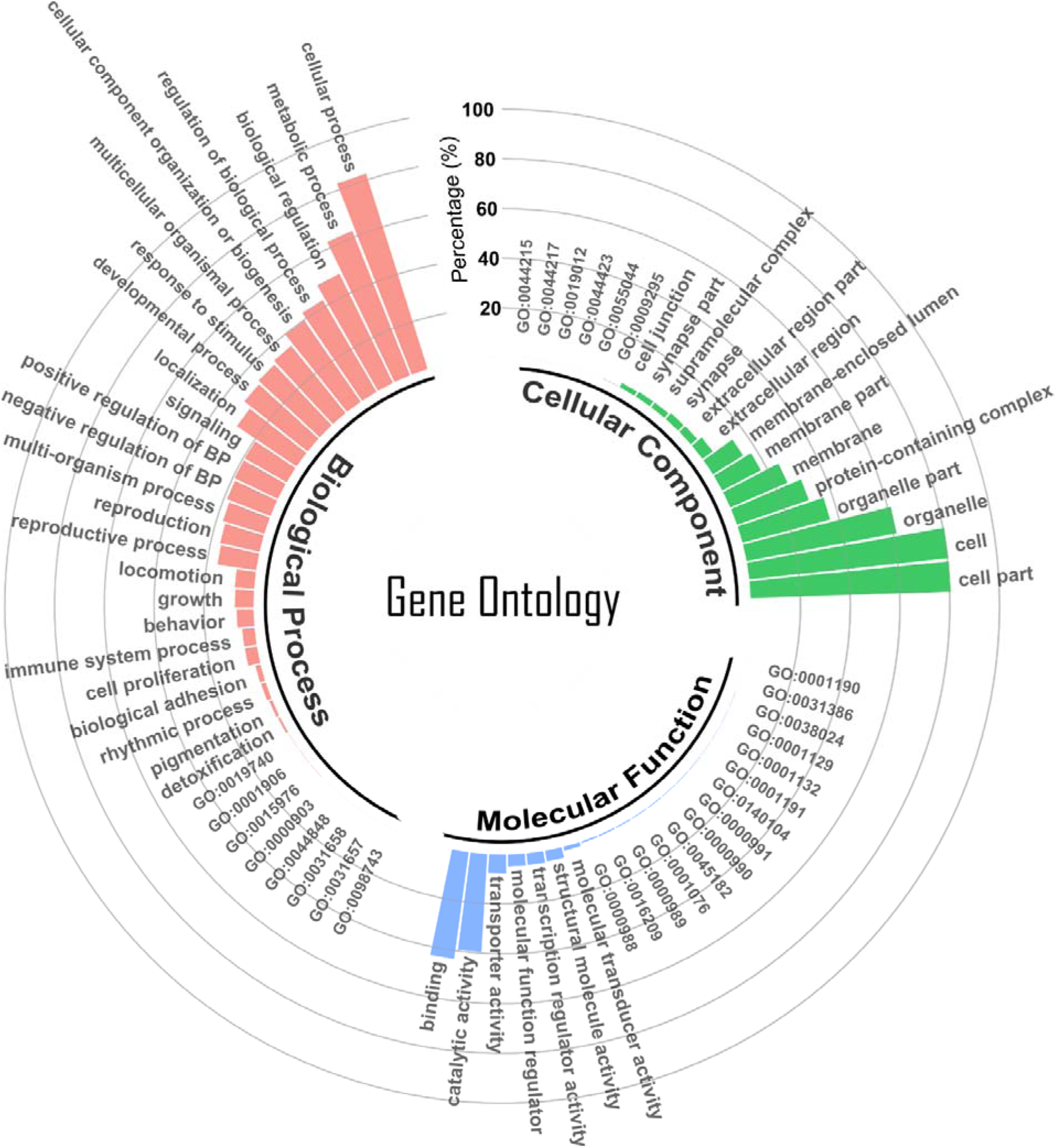
Gene Ontology classification of the *E. heros de novo* transcriptome.

Nearly 31,107 out of the 42,214 transcripts coding for predicted proteins with significant similarity to proteins available in the EggNOG database were classified into 24 COG functional categories (Fig. S1A). Predicted proteins in the *signal transduction* (3,532), *post-transcriptional modification, protein turnover, chaperone functions* (3,143), *transcription* (1,969), *carbohydrate metabolism and transport* (1,733), and *intracellular trafficking and secretion* (1,620) were among the most represented (Fig. S1A). A total of 21,248 sequences mapped to 411 KEGG pathways, with a high number of sequences mapping to *global and overview maps* (1,750), *signal transduction* (1,331), and *neurodegenerative diseases* (1,113) (Fig. S1B). *Global and overview maps* are intended to show global and overall features of grouped metabolism, with the *metabolic pathways* (894), *biosynthesis of secondary metabolites*(314) and *microbial metabolism in diverse environments* (138) pathways being the most represented (Table S2).

### 3.3. SNP identification

A total of 585,243 single nucleotide polymorphisms (SNPs) were detected, of which 173,672 SNPs survived the filtering protocols used. Transcript annotation indicated that most of the SNPs were detected in untranslated regions (UTRs), which were not further considered in this study. We identified 67,468 SNPs when comparing the susceptible and lambda-cyhalothrin-resistant strains (SUS vs PYR), with 9,588 non-synonymous SNPs that resulted in 4053 missense unigenes (Fig. 3 and Table S3).

**Fig. 3.**
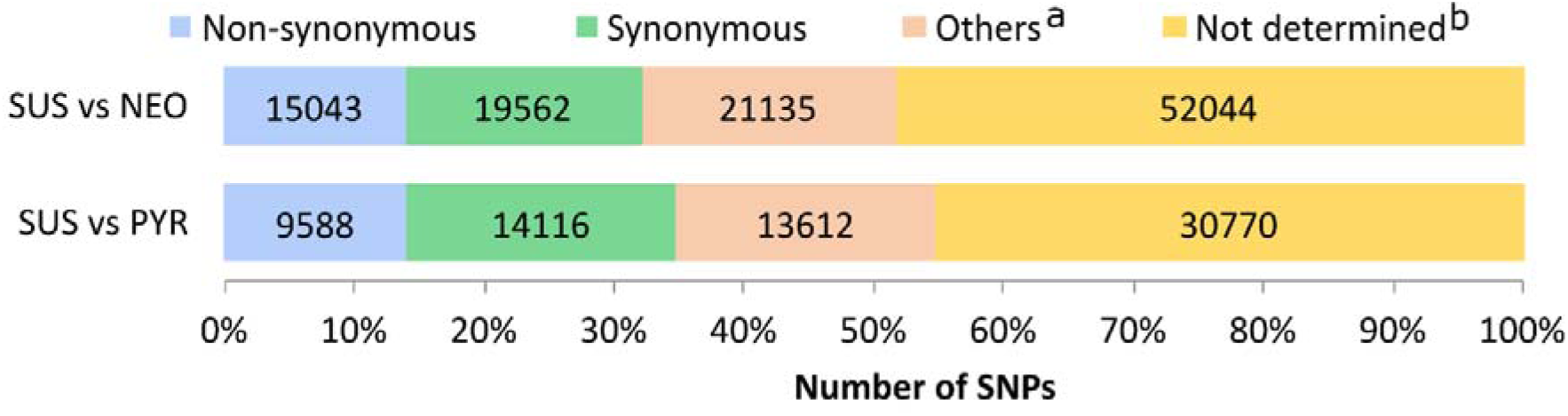
Annotation of SNPs by strain comparisons, SUS vs NEO and SUS vs PYR. **^a^** start codon gained or lost, stop codon gained or lost, 3’ UTR or 5’ UTR region. **^b^** SNP in a non-coding region of the transcript.

Comparisons of susceptible and thiamethoxam-resistant strains (SUS vs NEO) identified 55,740 SNPs in predicted protein-coding sequences, with 15,043 non-synonymous SNPs that resulted in 5549 missense unigenes (Fig. 3 and Table S4). The thiamethoxam-resistant strain carried a non-synonymous SNP (G208A) in the neuronal acetylcholine receptor subunit alpha 5 (CHRNA5), resulting in the amino acid substitution D70N in the transcript sequence assembled (DN8119_c0_g1_i2, Table S4). The prediction of transmembrane helices (TMHs) indicates this protein has four TMHs, and the D70N mutation was localized externally to the membrane (Fig. S2). Alignments of the extracellular domain of the neuronal acetylcholine receptor subunit alpha 5 from vertebrate and invertebrate species recognized several conserved residues, but the residues at position 70 of our contig was quite variable in all other species demonstrating that this is not a conserved residue (Fig. S2).

We identified missense variants with strong involvement in *metabolic and cellular process*, *gene expression regulation*, *signaling process*, *response to stress, transmembrane transport,* and *process of metabolization and excretion of xenobiotics* in both resistant strains (Fig. S3 and S4). Variants were identified in the *G-protein-coupled receptor* (GPCR) *methuselah* (*mth*), in *multidrug resistance-associated proteins, ATP-binding cassette subfamily* (ABC transporters) unigenes, and several unigenes annotated as enzymes acting in phase I and II of the detoxification process, such as *cytochrome P450*, *glutathione S-transferases*, *UDP-glucosyltransferases*, and *esterases* (Table S3 and S4).

### 3.4. Differential expression analysis

#### 3.4.1. Susceptible *versus* thiamethoxam-resistant *E. heros*

Comparative analysis of gene expression between the susceptible (SUS) and thiamethoxam-resistant (NEO) strains of *E. heros* identified 215 DE transcripts (p-adj ≤ 0.05 and log_2_FC ≥ 2), of which 139 were up-regulated and 76 were down-regulated in the NEO strain (Fig. 4A, B and Table S5). One-hundred and seventy-one (79.5%) of the DE transcripts were successfully annotated against the NCBI *nr* database, 80 against GO, 79 against KEGG, and 131 against COG. DE transcripts with GO annotation (80) were assigned to 44 functional groups (23 in *biological processes*; 14 in *cellular components;* 7 in *molecular functions*) (Fig. 5A). *Cellular* (GO:0009987) and *metabolic processes* (GO:0008152) were the most represented GOs in *biological processes; cell part* (GO:0044464) and *cell* (GO: 0005623) in *cellular components*; and *binding* (GO: 0005488) and *catalytic activities* (GO: 0003824) in *molecular processes*. GO enrichment analysis successfully identified 5 GO terms in *biological processes* (*cell communication*, *signaling*, *signal transduction*, *cellular response to stimulus*, and *cell surface receptor signaling pathway*), but only one term was enriched in the *cellular component* (*cytoplasm*) (Fig. 5B). COG analysis classified the DE transcripts into 24 functional categories, with *signal transduction* and *lipid metabolism* as the most represented (Fig. 6A). Finally, the DE transcripts were assigned to 135 KEGG pathways, with the largest number of transcripts mapping to the *endocrine system, signal transduction*, and *transport and catabolism pathways* (Fig. 6B).

**Fig. 4.**
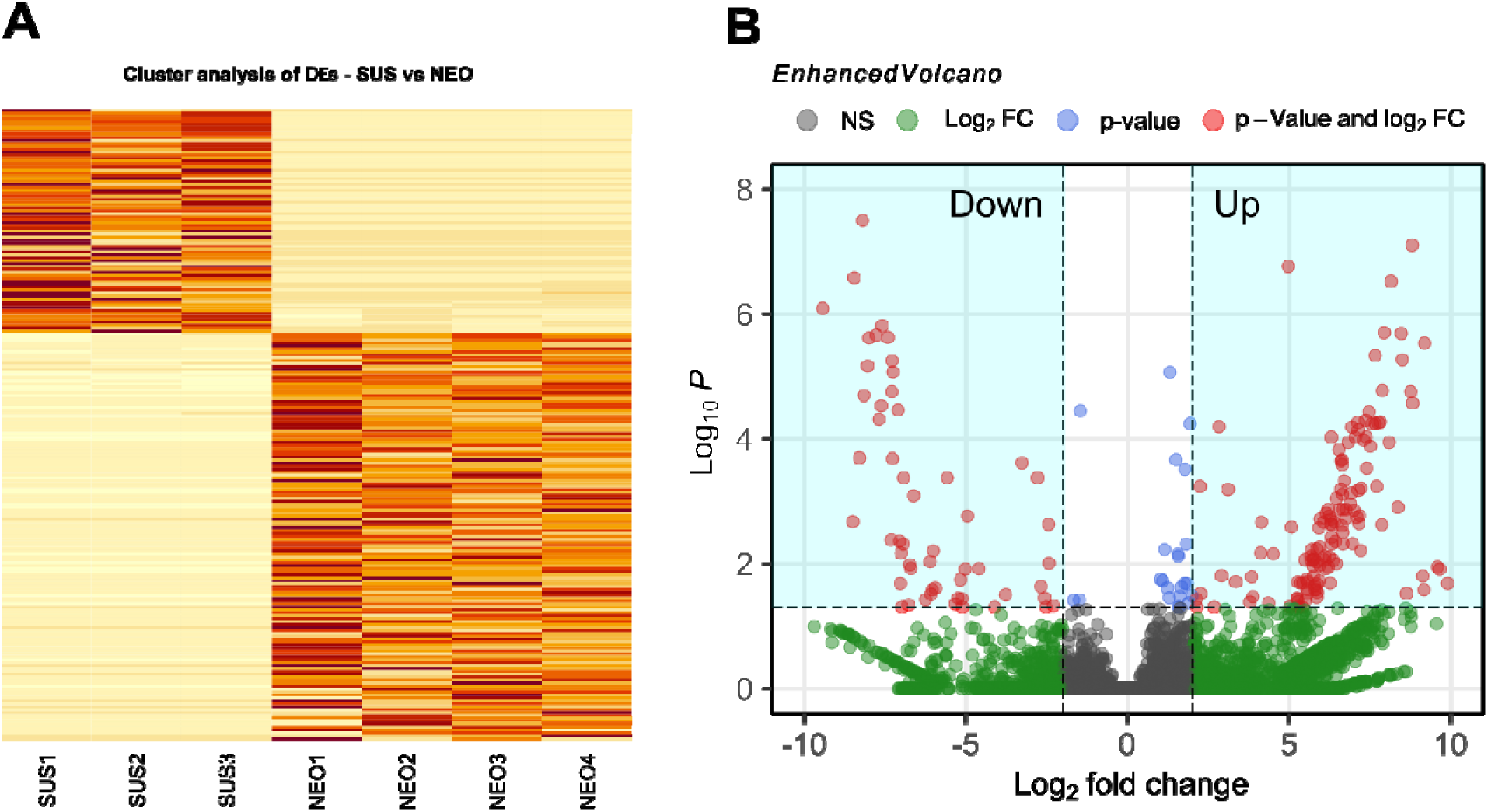
Heatmap and volcano plot of DEs between susceptible (SUS) and neonicotinoid-resistant (NEO) strains. **(A)** Heatmap of cluster analysis of DEs. The vertical axis represents the DEs, and the horizontal axis represents samples. **(B)** Volcano plot of DEs. The red dots represent up-regulated and down-regulated isoforms.

**Fig. 5.**
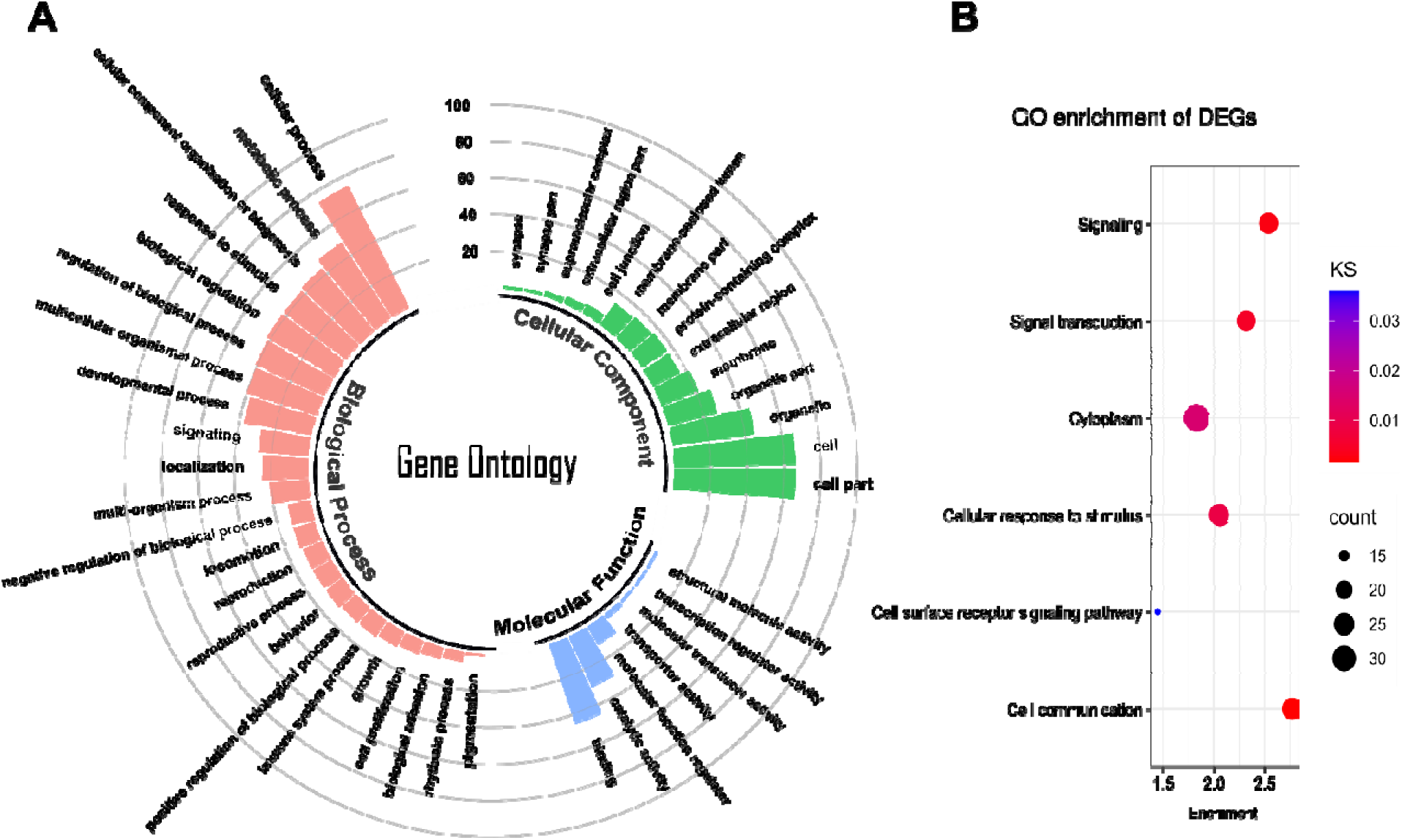
**(A)** Gene Ontology classification and **(B)** GO enrichment of DEs between neonicotinoid-resistant and susceptible strains of *E. heros*. KS (Kolmogorov-Smirnov, *p* < 0.05).

**Fig. 6.**
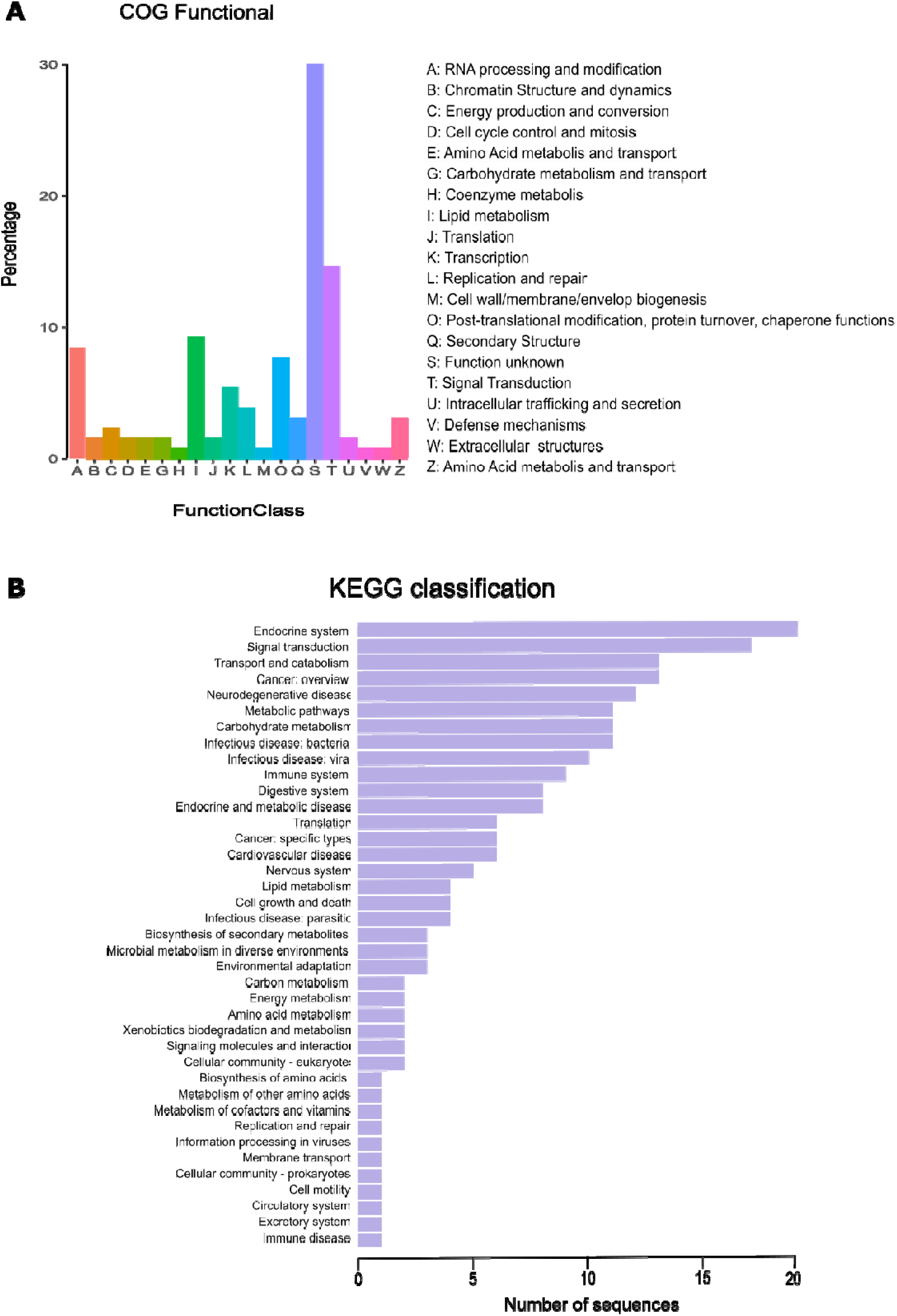
**(A)** Frequency of COG functional categories and **(B)** KEGG pathway classification of DEs between *E. heros* neonicotinoid-resistant and susceptible strains.

DE analysis identified the up-regulation of genes involved in signaling regulation mediated by G-protein coupled receptors (GPCRs), as the *secretin receptor-like* (DN5816_c0_g1_i9; log_2_FC = 6.64) and *cAMP-dependent protein kinase type I regulatory subunit* (DN850_c1_g1_i1; log_2_FC = 8.15) (Table S5). The DEs analysis also identified the up-regulation of transcripts related with detoxification of xenobiotics that are associated with insecticide resistance: two enzymes involved in phase II (conjugation) of the detoxification process, *glutathione-S-transferase* (DN935_c0_g1_i5; log_2_FC = 2.24) and a UDP-glycosyltransferase (DN1881_c0_g1_i5; log_2_FC = 2.83), and two translocases involved in phase III (transport), the *multidrug resistance protein 2* (DN14796_c1_g1_i5; log_2_FC = 5.57) and the *ABC transporter G family member 23-like* (DN10728_c0_g1_i4; log_2_FC = 8.09) (Table S5).

Insects have a complex immune system to defend themselves against micro and macroorganisms, and the DE immunity-related transcripts *polycomb group protein Psc-like* (DN11269_c0_g1_L1; log_2_FC = 6.60), *putative defensin* (DN129_c0_g1_i17; log_2_FC = 3.83), *ankyrin repeat domain-containing protein 17 isoform X11* (log_2_FC = 10.17) were up-regulated in the NEO *E. heros* strain. Several signal transduction transcripts were also up-regulated in NEO, such as the *guanine nucleotide-releasing factor 2 isoform X4* (log_2_FC = 7.30), *sterol regulatory element-binding protein 2* (log_2_FC = 7.04), *Ras-related protein Rap-2c* (log_2_FC = 8.76), *insulin-like growth factor-binding protein complex acid labile subunit* (log_2_FC = 7.09), and *wolframin-like isoform X2* (log_2_FC = 8.63) (Table S5).

#### 3.4.2. Susceptible *versus* lambda-cyhalothrin-resistant *E. heros*

We detected 204 DE transcripts (p-adj ≤ 0.05 and log2FC ≥ 2), with 106 up-regulated and 98 down-regulated transcripts in the PYR strain when compared to the SUS strain (Fig. 7A, B, Table S6). The functional annotation successfully annotated 171 DE (84%) (nr-NCBI), with 87, 88 and 134 been annotated against GO, KEGG and COG databases, respectively. Gene Ontology (GO) analysis assigned the functionally annotated transcripts into 44 functional groups and classified them into three categories: *molecular functions* (8), *biological processes* (24), and *cellular components* (12) (Fig. 8A). The main GO terms for *biological processes* were related with *cellular process* (GO:0009987) and *metabolic process* (GO:0008152); for the *molecular functions* were related to *catalytic activities* (GO: 0003824) and *binding* (GO: 0005488); and for *cellular components*, many annotated sequences were associated with *cell part* (GO:0044464) and *cell* (GO: 0005623). Eleven GO terms of *biological processes* and four of *cellular components* were enriched after GO enrichment analysis (KS, *p* < 0.05) (Fig. 8B). The COG analysis classified the DE transcripts into 20 functional categories. The functional categories *post-translational modification, protein turnover, chaperone functions*, *signal transduction*, and *lipid metabolism* were represented with frequencies greater than 10% (Fig. 9A). Analysis against the KEGG pathways database identified 161 pathways, with the largest number of transcripts represented in the *neurodegenerative disease* pathways, followed by *endocrine system*, *signal transduction*, and *transport and catabolism* (Fig. 9B).

**Fig. 7.**
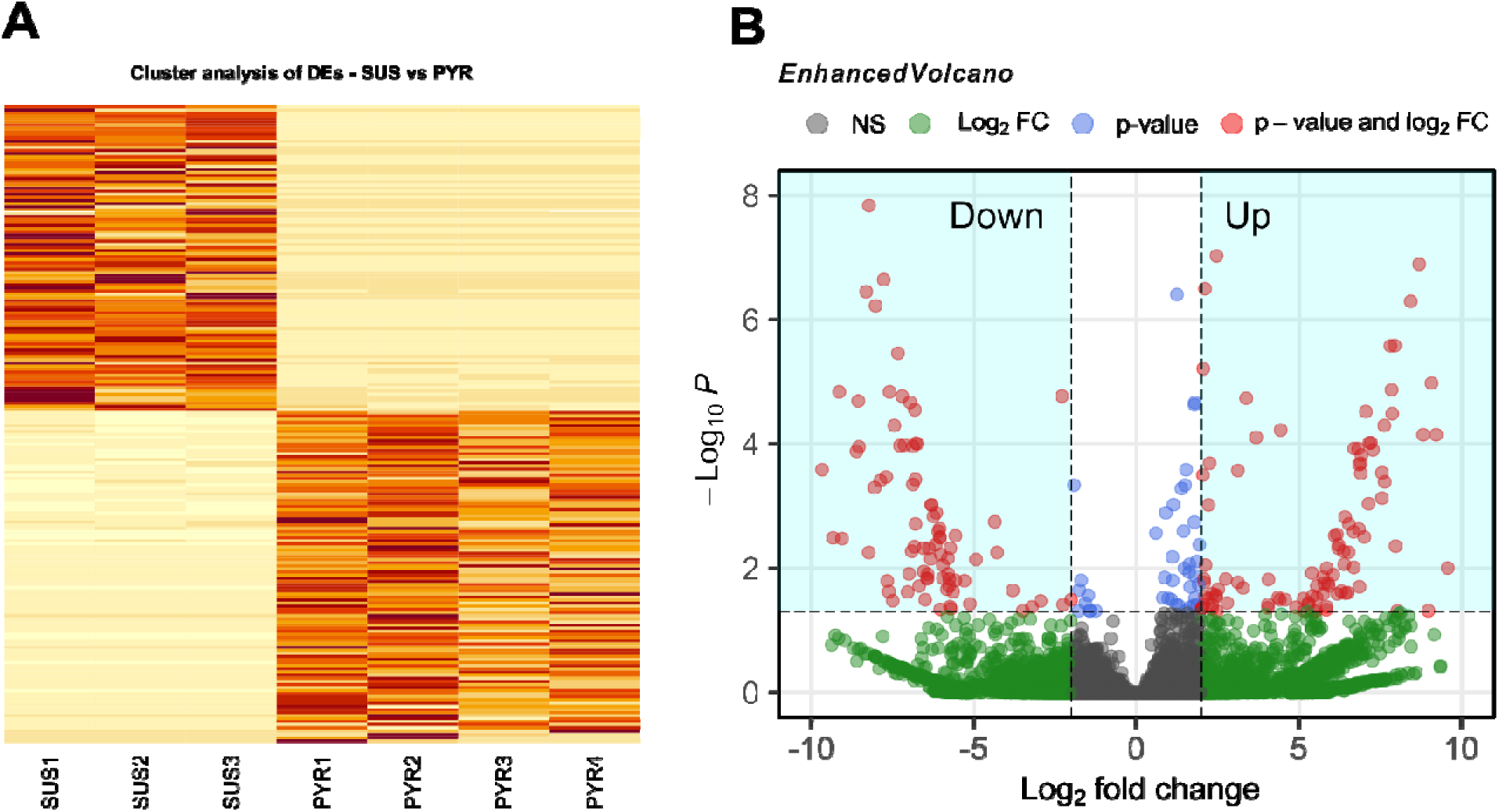
Heatmap and volcano plot of DEs between susceptible (SUS) and pyrethroid-resistant (PYR) strains. **(A)** Heatmap of cluster analysis of DEs. The vertical axis represents the DEs, and the horizontal axis represents samples. **(B)** Volcano plot of DEs. The red dots represent up-regulated and down-regulated isoforms.

**Fig. 8.**
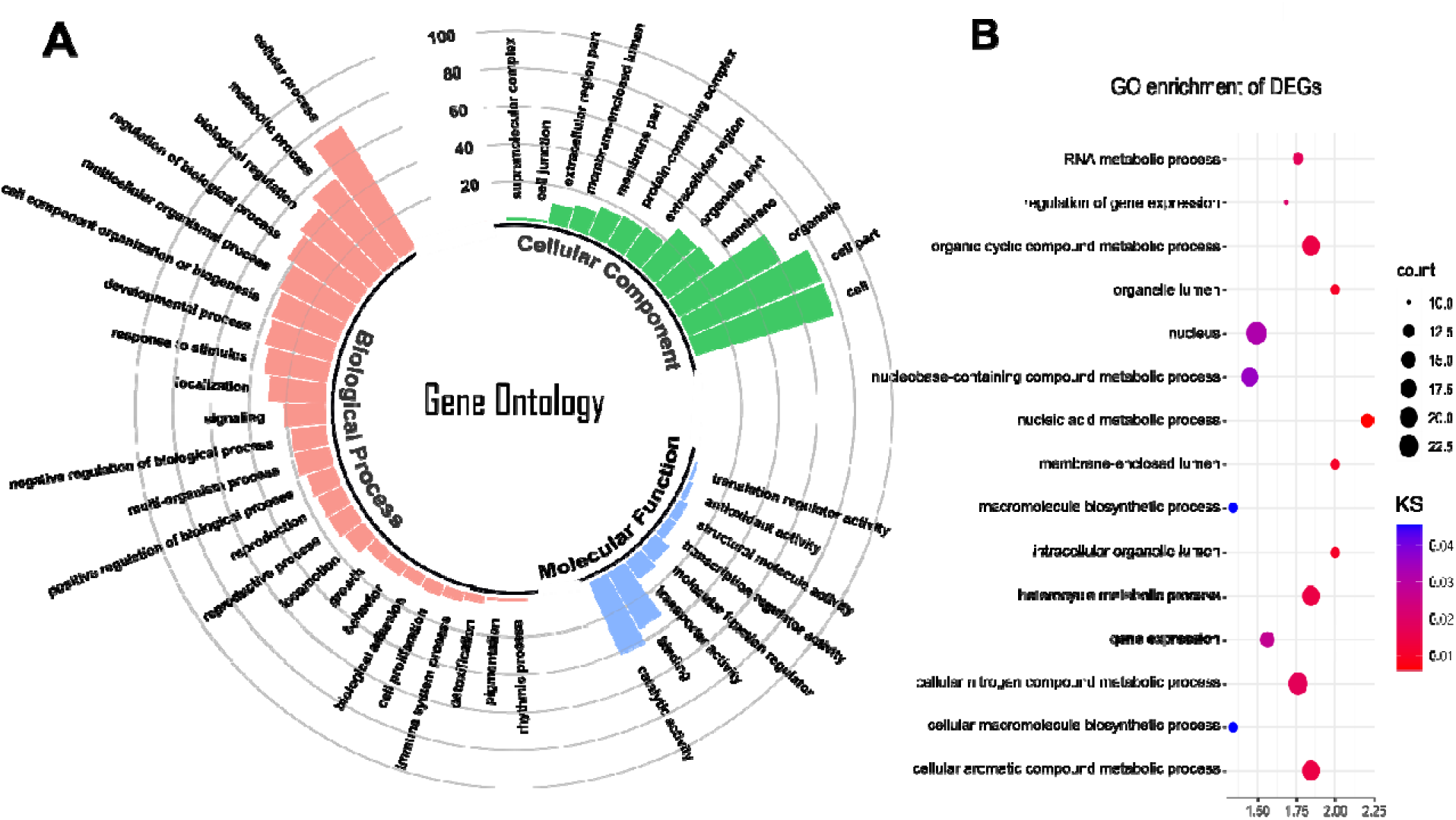
**(A)** Gene Ontology classification and **(B)** GO enrichment of DEs between pyrethroid-resistant and susceptible strains of *E. heros*. KS (Kolmogorov-Smirnov, p < 0.05).

**Fig. 9.**
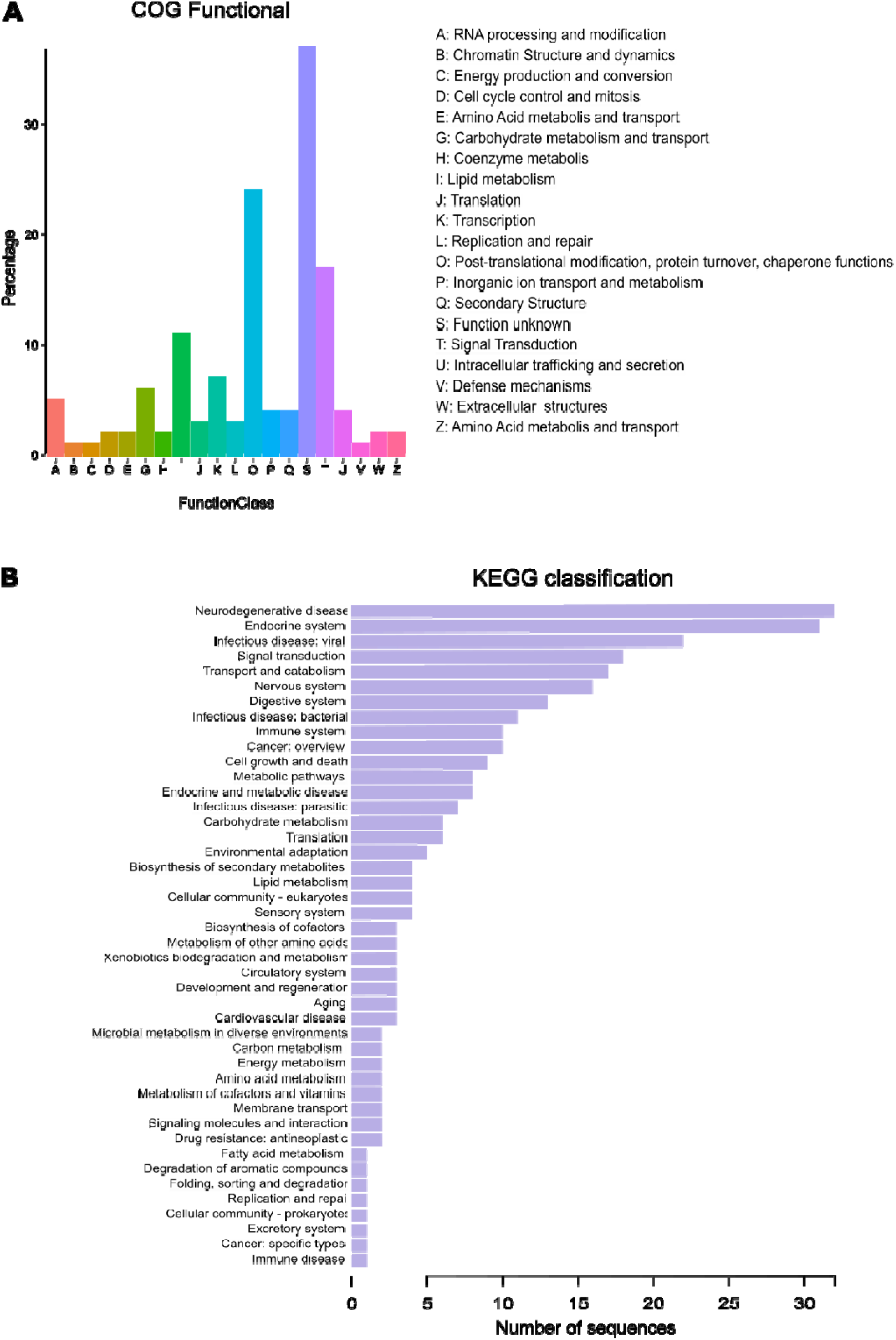
**(A)** Frequency of COG functional categories and **(B)** KEGG pathway classification of DEs between *E. heros* pyrethroid-resistant and susceptible strains.

DE analysis identified many DE transcripts related to critical life processes involved in insecticide resistance, such as the enhanced metabolism by detoxification and penetration resistance (Table S6). Several transcripts of enzymes and proteins involved in different steps of the process of detoxification were up-regulated in the PYR strain, such as the *esteraseFE4* (DN22_c1_g1_i12; log_2_FC = 2.45), *glutathione-S-transferase-like* (DN935_c0_g1_i2; log_2_FC = 2.46), UDP-glucosyltransferase (DN1881_c0_g1_i5; log_2_FC = 2.83), *multidrug resistance protein 1* (MRP1 = ABCC1) (DN2702_c1_g1_i1; log_2_FC = 2.05), and *ABC transporter G family member 23-like* (DN10728_c0_g1_i4; log_2_FC = 8.69). We also detected the up-regulation of the *larval cuticle protein A2B* (DN4787_c1_g1_i2; log_2_FC = 5.77) and *adult-specific cuticular protein ACP-20-like* (DN9039_c0_g1_i1; log_2_FC = 2.1) in the PYR strain (Table S6).

## 4. Discussion

Our data provide new insight into the putative molecular mechanisms underpinning resistance to pyrethroids and neonicotinoids in *E. heros*. Our transcriptomic analyses provide evidence that resistance in the NEO and PYR strains of *E. heros* is, at least in part, mediated by a metabolic component, represented by the overexpression of several proteins involved in the three phases of the metabolization of xenobiotics (Stanley, 2017; Stanley et al., 2009). The initial detoxification process was implicated by the discovery of the up-regulation of *esteraseFE4-like*, an enzyme acting in the functionalization of insecticides (phase I of the detoxification process) in PYR. Up-regulation of FE4 and E4 esterases were previously shown to be a widespread mechanism of insecticide resistance in the aphid *Myzus persicae* (Field and Devonshire, 1998; Tang et al., 2017), and other insects (Hsu et al., 2016; Zhu et al., 2004). The overproduction of esterases in *M. persicae* was initially reported to depend on gene amplification and gene methylation (Field et al., 1999). Later studies demonstrated the number of copies of the FE4/E4 genes can range from 6 to 104 in *M. persicae*, with a high correlation between esterase activity and the gene copy number. Esterase genes in aphids with low copy number were almost completely methylated, but no correlation was found between the gene methylation state and gene copy number (Bizzaro et al., 2005; Field et al., 1999).

A mechanism of metabolic resistance was also supported in our study by the observed increased expression of *glutathione-S-transferase* and UDP-glucosyltransferase (UGT), enzymes that mediate conjugation reactions (phase II) with insecticides, in the NEO and PYR strains. Increased activity of GSTs is commonly reported as a resistance mechanism to pesticides in insects (Huang et al., 1998; Vontas et al., 2002, 2001). UGTs are common, highly diverse enzymes that mediate the metabolization of xenobiotics in insects, including pyrethroids and neonicotinoids (Chen et al., 2023; Dimunová et al., 2022; Israni et al., 2022; Zhou et al., 2019). The specific UDP-glucosyltransferase overexpressed in our study was highly similar to the *2-hydroxyacylsphingosine 1-beta-galactosyltransferase-like* gene from *Halyomorpha halys. 2-hydroxyacylsphingosine 1-beta-galactosyltransferase* are UDP-glucosyltransferases involved in the catalytic reaction leading to the sphingolipid galactosylceramide. Galactosylceramides are cerebrosides commonly associated with vertebrates, while ceramide phosphoethanolamine is the cerebroside reported as the main constituent of myelinic membranes in invertebrates (Poitelon et al., 2020; Vacaru et al., 2013). Additionally, the domain families of *2-hydroxyacylsphingosine 1-beta-galactosyltransferases* from hemipterans differs from those of human *2-hydroxyacylsphingosine 1-beta-galactosyltransferases* (data not shown), supporting the different role of this UGT in insects.

Proteins acting on phase II of insecticide metabolization are involved in increasing the hydrophilicity of xenobiotics to improve their elimination by the excretory system (phase III). The enhanced elimination of xenobiotics (phase III) in the NEO and PYR strains was evidenced by the overexpression of transcripts belonging to subfamilies C and G of the superfamily of ATP-binding cassette (ABC) transporters, which are involved in efflux of insecticides in insects (Gott et al., 2017; Li et al., 2020; Merzendorfer, 2014). ABC transporters are thought to be associated with pyrethroid resistance in several insects (Bariami et al., 2012; He et al., 2019; Pohl et al., 2012; Rault et al., 2019). The NEO and PYR strains overexpressed the same ABCG (*ABC transporter G family member 23-like* gene). However, in the case of ABCCs, ABCC1 (*multidrug resistance protein 1*) was overexpressed in the PYR strain, while ABCC2 (*multidrug resistance protein 2*) was overexpressed in the NEO strain. The expression levels or activities of drug transporters have been demonstrated to be drug/tissue dependent, with the expression and activities of genetic variants in these drug transporters leading to drastic alterations on pharmacokinetics and drug response in a genotype-dependent manner (Bruckmueller and Cascorbi, 2021; Dermauw and Van Leeuwen, 2014). The increase in activity of efflux protein transporters suggests the requirement of higher diuresis for toxin clearance by the excretory system of insects for later excretion. Consequently, we might then expect to see increased activity of genes involved in diuresis, such as diuretic factors and aquaporins. There are several molecules that can stimulate diuresis in insects (diuretic hormone, kinin, serotonin, corticotropin releasing factor, and others), and they are essential for water balance and regulation of fluid secretion by Malpighian tubules in insects (Nation, 2022). We detected the up-regulation of a *diuretic hormone class 2* transcript (DN3531_c0_g1_i1) in both the NEO (log_2_FC =1.6; p-value = 0.06; p-adj = 0.99) and PYR (log_2_FC =2.2; *p-value* = 0.03; p-adj = 0.75) strains, however, in both cases we could not confirm their differential expression using the cut-off parameters set in our RNASeq analysis. We did not detect any variation in the level of expression of other insect osmoregulatory endocrine factors (kinin and serotonin) or in their receptors (serotonin receptor, tachykinin receptor, dopamine receptor, neuropeptide CAPA receptor, tyramine receptor, among others) observed in the transcriptome of *E. heros*.

Aquaporins are proteins involved in water transport in different tissues, and those in Malphigian tubules regulate water movement through the epithelium for urine formation (Nation, 2022; Spring et al., 2009). A DE aquaporin was identified as overexpressed in the PYR strain (DN2785_c0_g1_i3; log_2_FC = 8.8; p-value = 7.5e-08; p-adj = 7.1e-05), that was putatively identified as an *aquaporin AQPcic isoform X2* earlier reported in the filter chambers of the hemipteran *Cicadella viridis* (Caherec et al., 1996). This same aquaporin transcript was observed with a similar level of transcription in NEO (log_2_FC = 8.8; p-value = 0.0004), but with an adjusted p-value (p-adj = 0.07) above the cut-off originally set for the analysis (p-adj < 0.05). The DE aquaporin transcript observed in PYR is identical to the EUSHE_CONTIG_DN260927_c2_g2_i1 identified in a tissue-specific expression atlas for *E. heros* (BugAtlas, www.bugatlas.org), which is reported to be highly expressed in several tissues, including the gut and Malpighian tubules. The participation of aquaporin in the mechanism of insect resistance to insecticides has been reported in several studies (Mitchell et al., 2012; Tan and Chen, 2023), supporting the participation of the detoxification machinery as a mechanism of resistance, at least for the PYR strain.

The overexpression of two cuticle proteins containing the chitin-binding type R&R domain (CPR), *larval cuticle protein A2B* (DN4787_c1_g1_i2; log_2_FC = 5.77; p-adj = 0.01) and *adult-specific cuticular protein ACP-20-like* (DN9039_c0_g1_i1; log_2_FC = 2.1; p-adj = 0.02) in the PYR strain suggests that modifications in the cuticle could also contribute to *E. heros* resistance to lambda-cyhalothrin. The overexpression of CPRs alter the cuticle thickness and/or composition, slowing cuticular penetration of insecticides (Balabanidou et al., 2018; Fang et al., 2015; Huang et al., 2018).

Neonicotinoids bind to nicotinic acetylcholine receptors (nAChR) (Jones and Sattelle, 2010; Matsuda et al., 2020), but the α5 subunit of the nicotinic acetylcholine receptor is not recognized as a binding site of acetylcholine. The α5 and the β3 subunits are reported to have more of an accessory role in the receptor (Maskos, 2020). Mutations in the nAChR of neonicotinoid-resistant strains were always reported to occur in the subunits listed as binding sites of acetylcholine. Such mutations were reported in the α1/β2 subunits in *Drosophila melanogaster* (Perry et al., 2008), α1/α3 subunits (Y151Z) in *Nilaparvata lugens* (Liu et al., 2006, 2005), and in the β1 subunit in *Myzus persicae* (V101I and R81T) (Bass et al., 2011; Xu et al., 2022) and *Aphis gossypii* (R81T) (Hirata et al., 2017). Thus, it is unlike that thiamethoxam will bind to the α5 subunit. Moreover, multiple alignments of the extracellular domain of the α5 subunit of nAChR demonstrated the position where the D70N SNP was identified in NEO is not conserved. Thus, despite several mutations in nAChR have already been reported as target-site mutation mechanisms of neonicotinoid resistance in insects and that target-site mutations involved with neonicotinoid resistance in insects have been reported in different subunits of the nAChR, we do not believe the D70N SNP identified is involved in thiamethoxam resistance in the NEO strain of *E. heros*.

SNPs were also observed in *G-protein-coupled receptor* (GPCRs) *methuselah* (*mth*). GPCRs play important roles in physiological processes, toxicological response, and the development of insecticide resistance (Liu et al., 2021a, 2021b). *Methuselah* is a secretin-like receptor family involved in stress resistance and several other processes (Araújo et al., 2013; Brody and Cravchik, 2000), including insecticide resistance. *Musca domestica Mthl10* was reported as the resistance mechanism to the neonicotinoid imidacloprid (Ma et al., 2020) and *Lymantria dispar Ldmthl11* to the pyrethroid deltamethrin (Cao et al., 2019). SNPs variants were also detected in several other genes involved in resistance to insecticides, including enzymes acting in phase I and II, and transporter proteins of phase III of the detoxification process. These SNP variants require additional functional studies for the verification of their participation in the mechanism of resistance of the selected strains of *E. heros*.

In conclusion, our findings on thiamethoxam-and lambda-cyhalothrin resistant strains of *E. heros* implicate a shared metabolic resistance mechanism. Metabolic resistance in the lambda-cyhalothrin-resistant strain was also associated with a candidate mechanism of reduced penetration of insecticide through cuticle. In combination these findings provide a valuable list of candidate resistance genes and mutations for future functional investigation of their causal role in resistance. Such validated mechanisms will facilitate the development of diagnostics that can inform on the frequency and distribution of resistance in field populations of *E. heros* to aid rational control decisions and the development and implementation of resistance management strategies.

## Statements

### Author Contributions

Conceptualization: ECL, ARBN and FLC; Data curation: ECL; Formal analysis: ECL and ARBN; Funding acquisition: CO, CB and FLC; Investigation: ECL; Methodology: EL, ARBN and FLC; Project administration: CO and CB; Resources: CO provided *E. heros* strains; Supervision: FLC; Writing—original draft: ECL and FLC; Writing—review & editing: all authors read and approved the manuscript.

## Funding

Fundação de Amparo à Pesquisa do Estado de São Paulo (FAPESP) and the Biotechnology and Biological Sciences Research Council (BBSRC), UK BBSRC, UKRI provided a joint grant to CO and FLC (Fapesp grant# 2018/21155-6) and CB under the BBSRC-FAPESP Joint Pump-Priming Awards for AMR and Insect Pest Resistance in Livestock and Agriculture (Grant Ref: <u>BB/R022623/1</u> and <u>2017/50455–5</u>) and BBSRC-FAPESP Newton Award for AMR and insect pest resistance in agriculture and livestock (Grant Ref: <u>BB/S018719/1</u> and <u>2018/21155–6</u>). Fapesp also provided a post-doctoral fellowship to ARBN (grant# 2019/17215-6). This study was financed in part by the Coordenação de Aperfeiçoamento de Pessoal de Nível Superior - Brasil (CAPES) - Finance Code 001. CAPES provided a doctoral fellowship to ECL.

### Data Availability Statement

Illumina data will be publicly available at the NCBI BioProject PRJNA970298.

