## Supplementary material for "Transcriptomic investigation of the molecular mechanisms underlying resistance to the neonicotinoid thiamethoxam and the pyrethroid lambda-cyhalothrin in *Euschistus heros* (Hemiptera: Pentatomidae)": Table S: Supplementary material 1.pdf

Lira et al. (2023) - Supplementary Material Fig S1 - Fig S4.

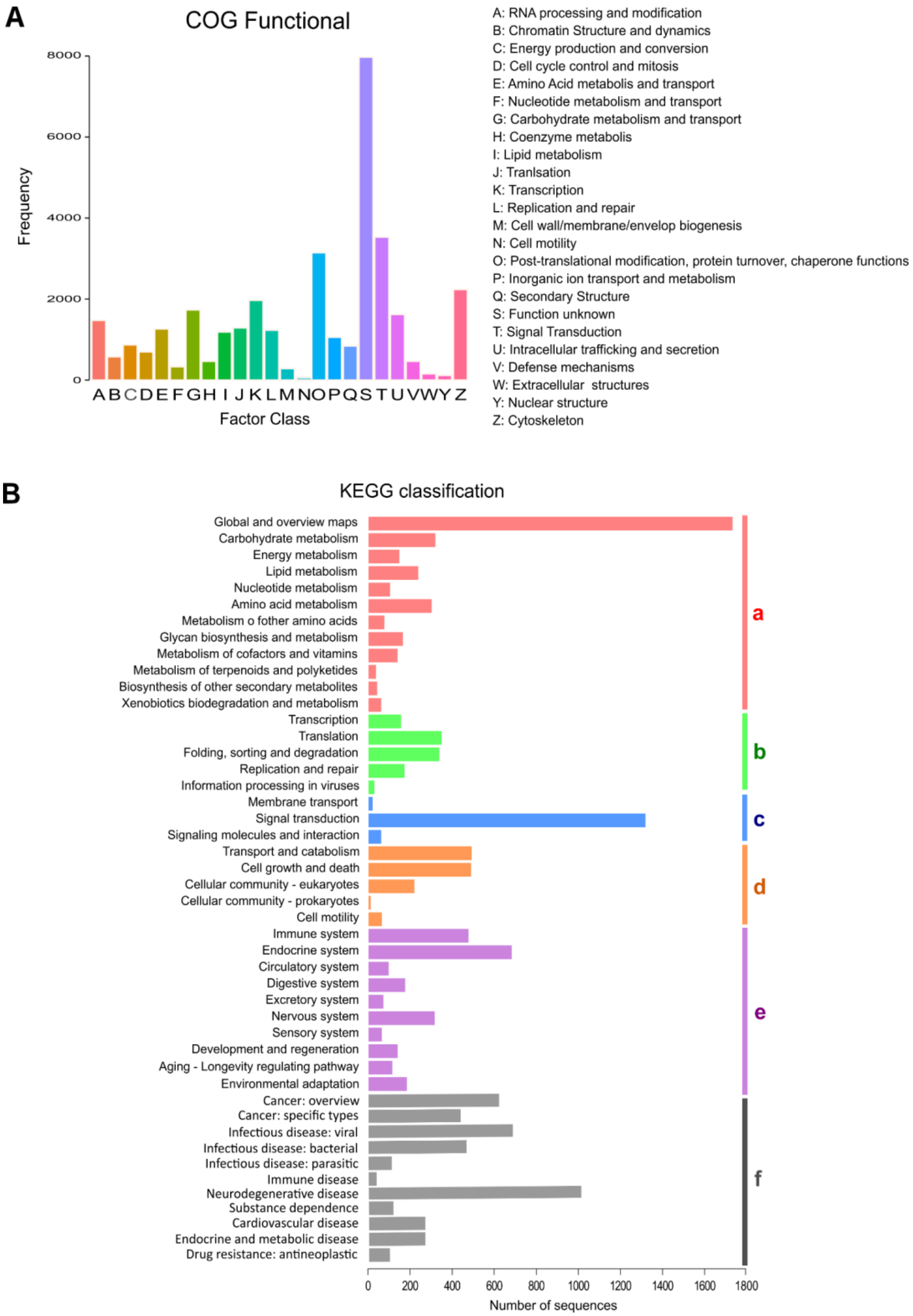


**Fig. S1. (A)** Frequency of COG functional categories and **(B)** KEGG pathway classification. Sequences were divided according to the pathways in which they participated: (a) Metabolism; (b) Genetic Information Processing; (c) Environmental Information Processing; (d) Cellular Processes; (e) Organismal Systems; (f) Human Disease.

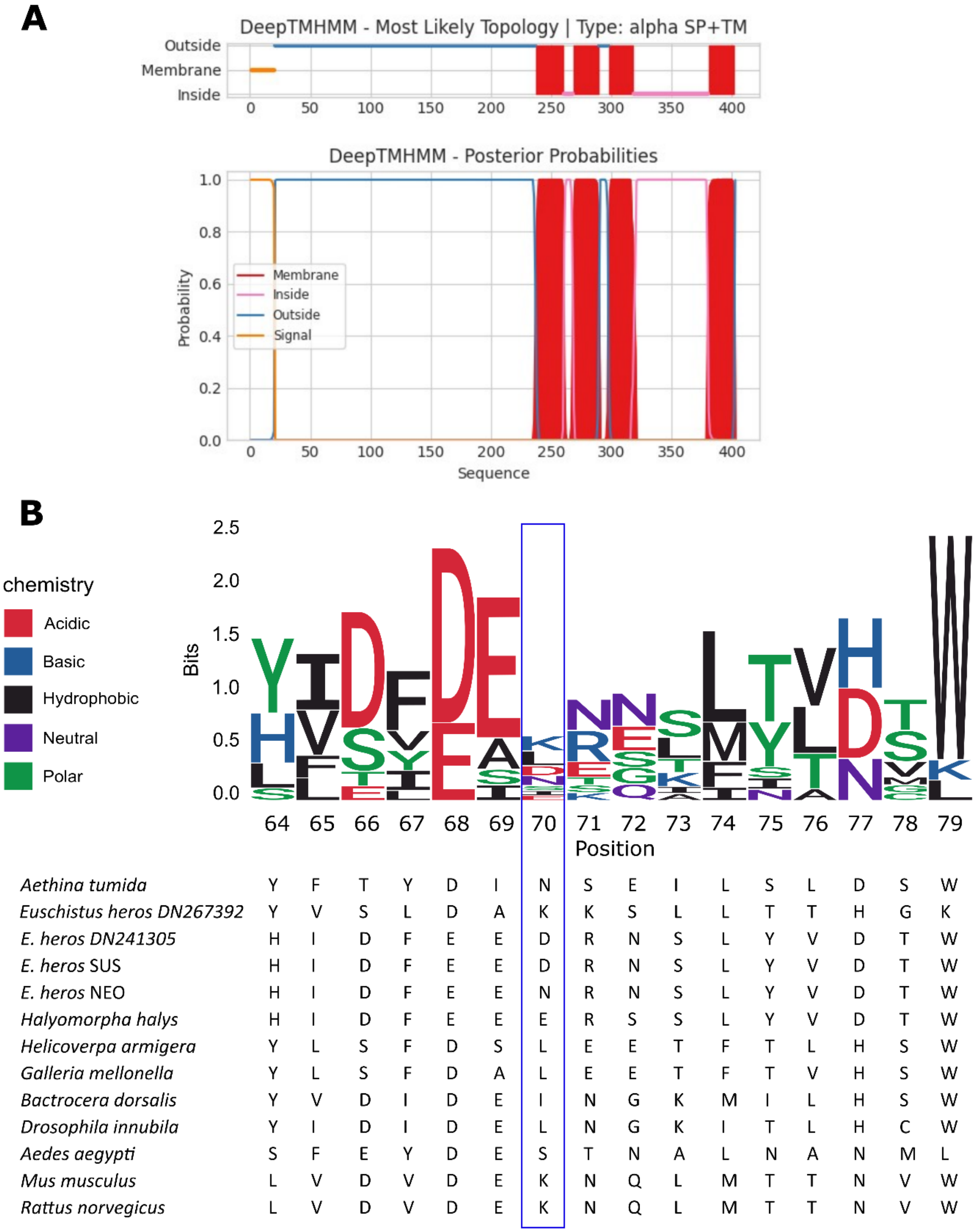


**Fig S2. (A)** Alpha and beta transmembrane proteins using deep neural networks of the neuronal acetylcholine receptor subunit alpha 5 (CHRNA5) of *E. heros* using DeepTMHMM and **(B)** ClustalW alignment of the amino acid sequence of a portion of the extracellular domain LGIC_ECD_Cation of the alpha 5 subunit of vertebrate and insect nicotinic acetylcholine receptors containing the D70N SNP detected in the NEO strain of *E. heros*. Consensus sequences of *E. heros* NEO and SUS strains were aligned against the LGIC_ECD_Cation domain of the CHRNA5 of *Aethina tumida* (XP_019865617.2), *E. heros* BugAtlas (DN267392 c0 g1 i2), *E. heros* BugAtlas (DN241305_c0_g1_i1), *Halyomorpha halys* (XP_014277544.1), *Helicoverpa armigera* (XP_021200871.2), *Galleria mellonella* (XP_026763934.1), *Bactrocera dorsalis* (JAC54180.1), *Drosophila innubila* (XP_034474756.1), *Aedes aegypti* (XP_001650053.1), *Mus musculus* (NP_001360830.1), and *Rattus norvegicus* (NP_058774.2).

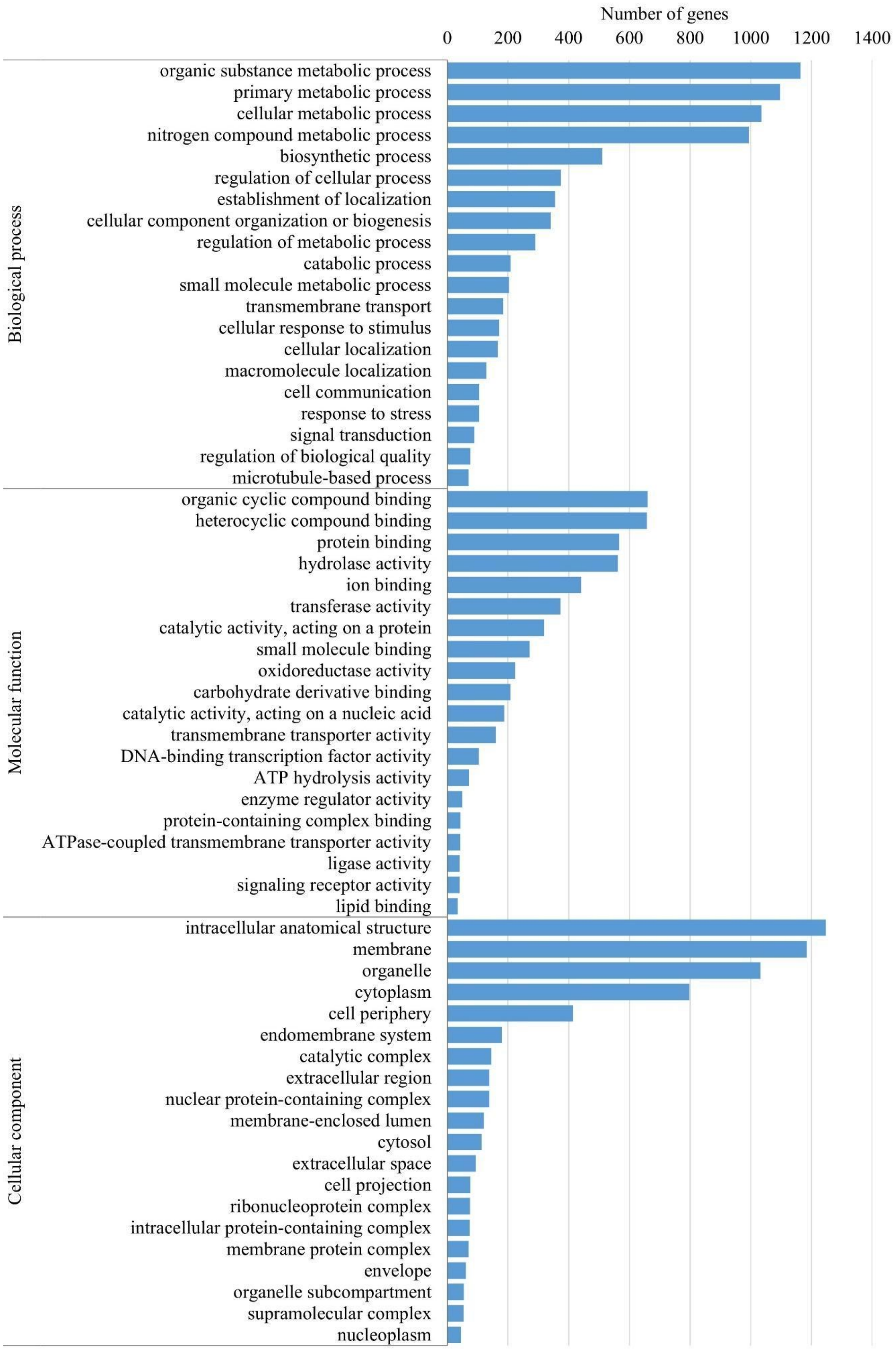


**Fig. S3**. Gene Ontology classification of unigenes with missense variants identified comparing susceptible and resistant *E. heros* strains to thiamethoxam (SUS vs NEO).

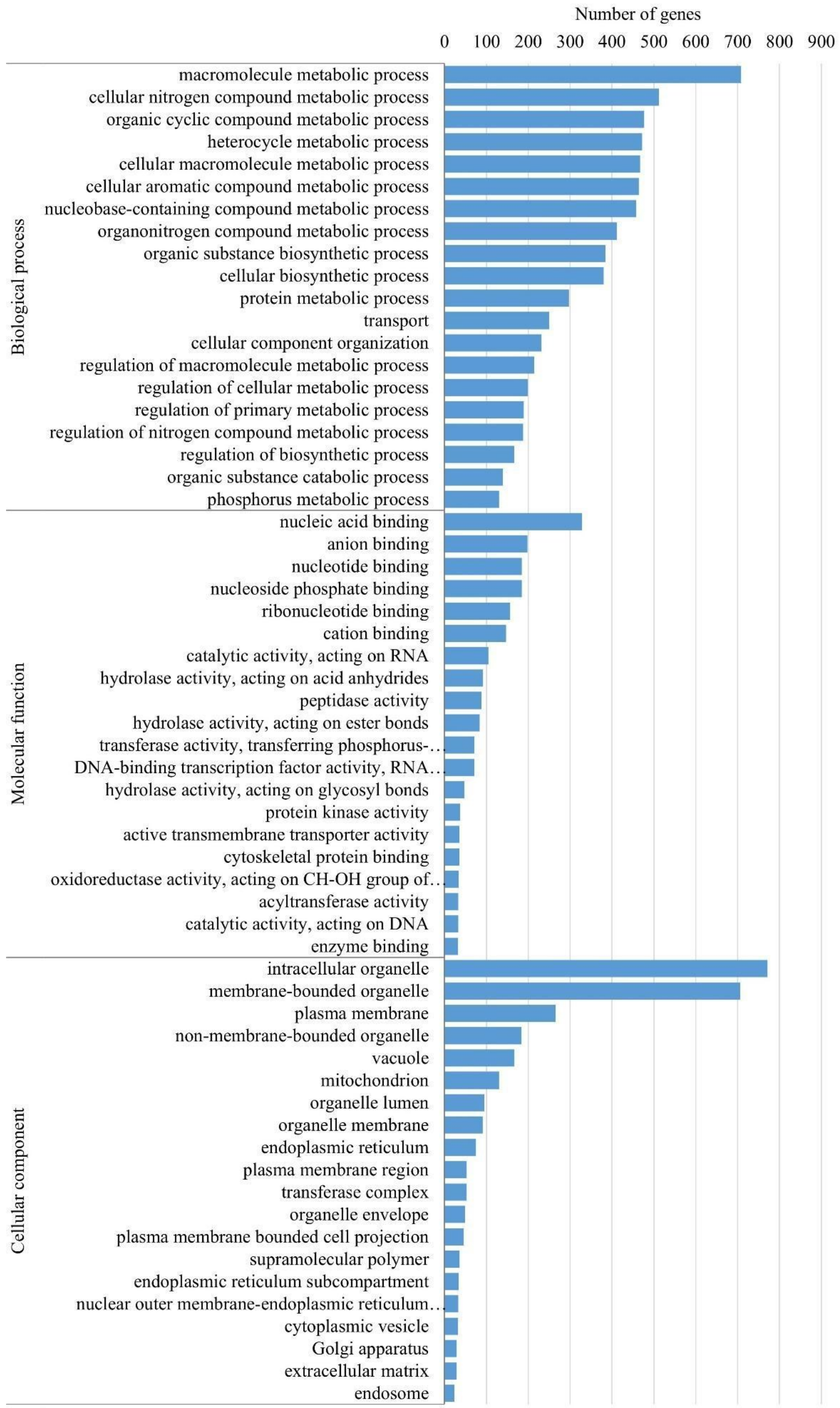


**Fig. S4**. Gene Ontology classification of unigenes with missense variants identified comparing susceptible and resistant *E. heros* strains to lambda-cyhalothrin (SUS vs PYR).
